# Sex-specific growth and lifespan effects of germline removal in the dioecious nematode *Caenorhabditis remanei*

**DOI:** 10.1101/2023.12.07.570570

**Authors:** Martin I Lind, Brian S Mautz, Hanne Carlsson, Andrea Hinas, Erik Gudmunds, Alexei A Maklakov

**Affiliations:** Animal Ecology, Department of Ecology and Genetics, Uppsala University, Uppsala, Sweden; Department of Environmental and Biosciences, Halmstad University, Halmstad, Sweden; Population Analytics & Insights, Data Sciences Analytics & Insights, Innovative Medicine Research & Development, Johnson & Johnson, Spring House, PA, USA; School of Biological Sciences, University of East Anglia, Norwich, UK; Department of Cell and Molecular Biology, Uppsala University, Uppsala, Sweden; Evolutionary Biology, Department of Ecology and Genetics, Uppsala University, Uppsala, Sweden

**Keywords:** Germline, glp-, Heat-shock, Lifespan, Sex-specific

## Abstract

Germline regulates the expression of life-history traits and mediates the trade-off between reproduction and somatic maintenance. However, germline maintenance in itself can be costly, and the costs can vary between the sexes depending on the number of gametes produced across the lifetime. We tested this directly by germline ablation using glp-1 RNAi in a dioecious nematode *Caenorhabditis remanei*. Germline removal strongly increased heat-shock resistance in both sexes, thus confirming the role of the germline in regulating somatic maintenance. However, germline removal resulted in increased lifespan only in males. High costs of mating strongly reduced lifespan in both sexes and obliterated the survival benefit of germline-less males even though neither sex produced any offspring. Furthermore, germline removal reduced male growth before maturation but not in adulthood, while female growth rate was reduced both before and especially after maturation. Thus, germline removal improves male lifespan without major growth costs, while germline-less females grow slower and do not live longer than reproductively functional counterparts in the absence of environmental stress. Overall, these results suggest that germline maintenance is costlier for males than for females.

## Introduction

Ageing is the physiological deterioration of organismal function and increases the likelihood of death with age, thus limiting lifespan ^1^. Since the strength of selection is reduced with advancing age ^2^, evolutionary theory of ageing postulates that ageing evolves as a result of selection for genes with beneficial effects early in life, despite late-life costs ^3^ and accumulation of mutations with late-life effects ^4^. Since all cells are subject to wear and tear, long lifespan requires effective repair mechanisms, to perform somatic maintenance ^5^. High levels of maintenance can be costly, and evolution of long life is generally associated with trade-offs such as reduced reproduction, slow early life development and/or growth ^6–8^.

Perhaps the most well-known trade-off is that between reproduction and lifespan – the so-called ‘cost of reproduction’ ^9^. Traditionally, the major cost of reproduction in species without parental care has been seen as the direct cost of producing gametes ^10^. However, gamete production is not the only costly expenditure associated with the germline. While the soma is disposable, the germline stem cells are essentially immortal and germline is maintained across generations ^5^. This immortality should come with a cost of germline maintenance and repair ^11^. Indeed, several studies suggest that germline maintenance is costly ^12,13^. For example, in *Caenorhabditis elegans*, there is a trade-off between somatic and germline maintenance ^14^ and low-condition (low quality) *Drosophila melanogaster* flies have more germline mutations, potentially because low-condition individuals invest less in high-fidelity repair ^15^. Moreover, not only the germline genome but also its proteome seems to be protected to a higher degree than the somatic proteome ^16,17^.

In theory, if the germline is experimentally removed, resources freed up from germline maintenance, could be used for increased somatic maintenance, and as a result, increased lifespan. In line with this hypothesis, germline removal in *Drosophila* ^18^ or *C. elegans* hermaphrodites ^19–21^ is associated with increased lifespan. However, while early studies suggested that lifespan-extending effect of germline removal is evolutionarily conserved, germline ablation is not universally associated with lifespan extension across taxa, and instead can lead to increased growth ^22^. Similarly, germline removal resulted in increased somatic repair under stress in male zebrafish leading to enhanced re-growth of lost fin tissue ^23^.

The cost of germline maintenance is however likely to differ between males and females ^11^. The two sexes are often defined by the relative size of their gametes; while females produce fewer and larger gametes, male gametes are small and plentiful. This anisogamy results in different selection pressures for the two sexes ^24^, and this sex-specific selection often results in differences in lifespan ^25^. However, the sexes may also differ in the amount of germline maintenance required across life course. Since males produce more gametes than females, they also need to maintain and repair a larger number of germline stem cells throughout their life. Since maintenance of the germline and repair of mutations seems to be expensive ^12,13,23^, males are likely to pay a larger maintenance cost than females. As a consequence, removal of the germline is expected to extend the lifespan of males more than that of females ^11^.

It is important to note that resource allocation is not the only and not necessarily even the main cause of life-history trade-offs. Reproduction and survival are shaped by cellular signalling networks. For example, the reproduction-lifespan trade-off is greatly influenced by the insulin/IGF-1(IIS) and mTOR signalling pathways ^26,27^. Genetic manipulations of these pathways have revealed that traits normally in trade-offs can be experimentally uncoupled ^28–30^. Additionally, germline-soma signalling determines the effect of germline removal on lifespan. In particular, the somatic gonad (the non-germline component of the gonad) signalling is key for lifespan-extending effects of germline removal. If the whole gonad (both somatic gonad and germline) is ablated in *C. elegans*, lifespan is unaffected. Only when ablating the germline precursor cells, leaving the somatic gonad intact, is lifespan extended^19^. According to one model, the functioning germline blocks the somatic signal, which enables reproduction at the cost of reduced somatic maintenance ^21^. Following this logic, the lack of a germline mimics a non-proliferating germline, which enables the soma to carry out maintenance, resulting in extended lifespan ^31,32^. While this suggests that germline-soma signalling is key for lifespan extension, these experiments do not directly discount the potential for resource allocation trade-offs. It is possible that the presence of functional germline syphons resources into germline maintenance, away from somatic maintenance, as suggested by the ‘expensive germline’ hypothesis ^11,23^.

We set out to test the sex-specific effects of germline removal on stress resistance, development time, growth, body size and survival in the dioecious nematode *Caenorhabditis remanei*, a sister species to the well-known model organism *C. elegans*. Germline-less worms were produced using *glp-1* RNAi. GLP-1 is a receptor in the LIN-12/Notch family which mediates the mitosis/meiosis decision in the *C. elegans* germline, and is therefore vital for germ line proliferation ^33,34^. GLP-1 is conserved across *Caenorhabditis* nematodes, including *C. remanei* ^35^. Most loss-of function mutations in the *glp-1* gene make germ cells that would otherwise divide mitotically to instead go into meiosis, and as a result, in *C. elegans*, only a few sperms are produced, and no eggs ^33^. We found that germline removal increases stress resistance in both sexes, but the effects on growth and lifespan were highly sex-specific, suggesting that the germline regulates investment in life-history traits differently in males and females.

## Methodology

### Worm maintenance

We used the wild-type SP8 strain of *C. remanei*, obtained from N. Timmermeyer at University of Tübingen. This is a cross of three other strains, and harbour substantial genetic variation (e.g. ^36,37^). Worms were maintained under standard laboratory conditions in dark incubators at 20°C on nematode growth medium (NGM) agar plates, following standard procedures ^38^.

### RNAi

RNAi against *C. remanei glp-1* was carried out by microinjection, essentially as described previously ^35^. First, total RNA was prepared from mixed stage *C. remanei* using TRIzol (ThermoFisher Scientific) and mechanical disintegration with a syringe with a 18-20 gauge needle. After chloroform extraction, RNA was precipitated by addition of isopropanol, washed with 75 % ethanol and dissolved in water. DNase I (ThermoFisher Scientific) digestion was performed according to the manufacturer’s instructions, followed by phenol/chloroform/isoamyl alcohol (25:24:1) extraction and ethanol precipitation. Subsequently, cDNA was prepared by reverse transcription using an oligo(dT)_18_ primer and RevertAid H Minus reverse transcriptase (ThermoFisher Scientific) according to the manufacturer’s instructions.

*glp-1* was PCR amplified from cDNA using Phusion DNA polymerase (ThermoFisher Scientific) and primers CR41 and CR42 ^35^, adding T7 promoters on each end. Cycling conditions were 98°C for 30 s, followed by 40 cycles of 98°C for 10 s, 55°C for 30 s and 72°C for 30 s, and a final extension of 1 min. DNA encoding green fluorescent protein (*gfp*) to be used as non-target control for RNAi injections was amplified from plasmid pPD95.75 using primers 241_GFP_forw_pPD95.75 (5’-TAATACGACTCACTATAGGGCAAATTTTCTGTCAGTGGAG-3’) and 242_GFP_rev_pPD95.75 (5’-TAATACGACTCACTATAGGGGTTACAAACTCAAGAAGGACC-3’) with Phusion DNA polymerase using the same cycling conditions as for *glp-1* but with 35 cycles.

The *glp-1* and *gfp* PCR products were used as templates for in vitro transcription using T7 RNA polymerase (Life Technologies) according to the manufacturer’s instructions. RNA integrity was determined by agarose gel electrophoresis. In vitro transcribed dsRNA (1 mg/ml) was injected into mated *C. remanei* females one day post L4 and progeny of injected mothers were DAPI stained at L4 stage as described previously ^39^.

One hour after injection, worms were moved individually to new plates, where they were allowed to lay eggs for five hours (one hour for the development and size assays). Mothers were then kept on separate plates, where we monitored non-hatching eggs two days later. Offspring of mothers who laid hatching eggs after 5h were discarded.

Eggs from the egg-laying plates were allowed to develop until L4, when they were isolated and used in the assays below (except for development and size assays, where they were followed from egg). Control worms (injected with *gfp*) were treated the in same manner.

Germline removal using *glp-1* RNAi completely abolished reproduction when worms were kept in mixed-sex groups.

### Heat shock assay

Heat shock survival assays were performed on mated worms at day 2 of adulthood. Worms, in total 50 per sex and treatment, were placed in sex-specific groups of 10 worms on seeded 35 mm NGM plates and placed in a climate chamber set to 40°C. Since males are more heat shock resistant than females ^37^, we ran separate assays for the two sexes. Therefore, females were subjected to heat-shock for 90 minutes, while males were exposed for 95 minutes. After the heat-shock, worms were returned to 20°C. Heat shock survival was scored 24h after the heat-shock treatment, and worms were considered dead if they did not respond to touch and showed no signs of pharyngeal pumping.

### Lifespan assay

Lifespan assays for virgin worms were initiated by placing individual worms in late L4 stage on individual 35 mm NGM plates (n = 100 for each sex and treatment). Worms were then transferred daily to new plates. Worms were scored as dead if they did not respond to touch and pharyngal pumping had ceased. Missing worms and females dying of matricide (internal hatching of eggs) were censored. For the mated treatment (n = 50 for each sex and RNAi combination), worms were kept in company with a control individual from the opposite sex. Because of high infection rate of unwanted bacteria in the mated experiments, worms from infected plates were moved to plates containing the antibiotic kanamycin for the rest of the experiment.

### Development time and size assay

Newly injected worms were allowed to lay eggs for 1h. From 59 h after egg laying, plates were checked every hour, and sexually mature females (26 control and 15 *glp-1*) and males (19 control and 14 *glp-1*) were scored for development time to the nearest hour and photographed using a Lumenera Infinity2-5C digital microscope camera mounted on a Leica M165C stereo microscope. After photography, mature worms were kept individually on 35 mm NGM plates and were moved to new plates each day. Four days after maturity, all individuals were photographed again (1 control female, 7 control males and 3 *glp-1* males were lost before day 4). Since female size peak at day four, and male size has reached its asymptote ^40^, this day was chosen to represent adult size. Size was estimated as cross-section area (in mm^2^) and measured using the image analysis program *ImageJ* (https://imagej.nih.gov/ij/).

### Statistical analyses

Heat shock survival was analysed as linear models with a binomial error distribution, where each plate (with 10 individuals) was a replicate. RNAi treatment was modelled as a fixed factor, and since each plate consist of one family, no random effect was added.

Lifespan was analysed in Cox proportional hazard models with Gaussian random effects using the *coxme* package ^41^. RNAi treatment was fitted as a fixed effect, and family effects were accounted for by adding mother ID as a random effect. For the experiment with mated worms, we also added infection presence (yes, no) as a fixed factor, since plates with bacterial infection (which only occurred in these two experiments) were treated with the antibiotic kanamycin.

Development time, size at maturity and adult size were analysed in linear mixed-effect models implemented in the package *lme4* ^42^. RNAi treatment was treated as a fixed effect, and family (mother ID) was modelled as a random effect.

## Results

We found that germline removal increased heat shock resistance of both female (χ^2^ = 24.72, d.f. = 1, p < 0.001) and male (χ^2^ = 12.60, d.f. = 1, p < 0.001) worms (fig. 1).

**Figure 1.**
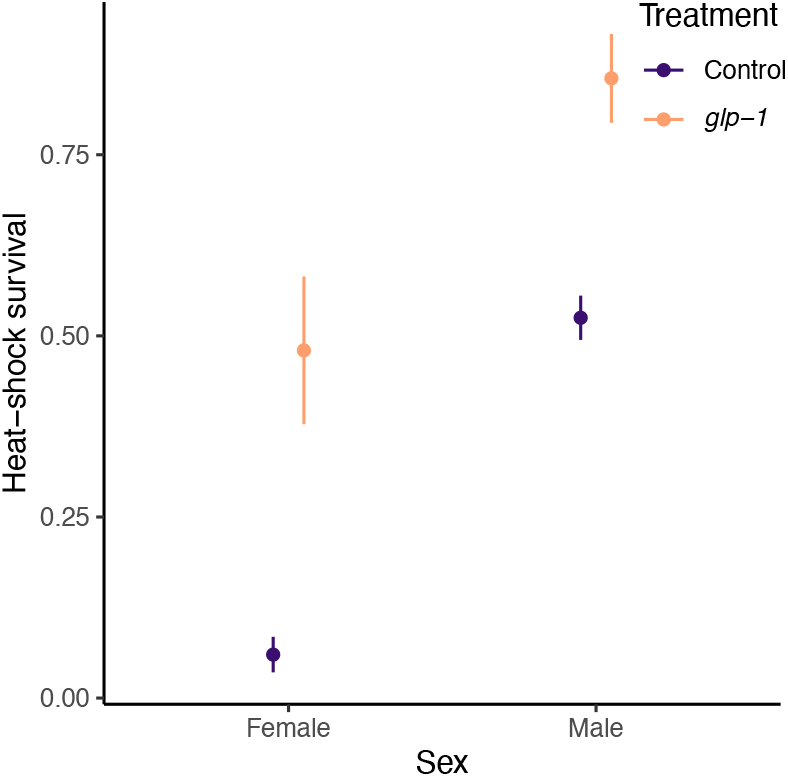
Sex-specific heat shock survival (proportion surviving) of control (purple) of germline-less *glp-1* treated (orange) worms. Error bars represent mean ± SE. Females were exposed to 40°C for 90 minutes, while the more heat-shock resistant males were exposed for 95 minutes, therefore the sexes should not be directly compared.

We also found a sex-specific lifespan effect for virgins (fig. 2). While germline removal did not influence lifespan of virgin females (z = -0.68, p = 0.490) it extended the lifespan of virgin males (z = -2.22, p = 0.027). In contrast, we did not find any lifespan effect of germline removal for mated worms (fig. S1), neither for females (z = 0.17, p = 0.870) nor for males (z = -0.07, p = 0.940). Interestingly, in the experiments on mated worms, 40 of 100 female plates and 68 of 100 male plates were infected by unwanted bacteria and we found a strong effect of bacterial infection and the subsequent use of the antibiotic kanamycin on these infected plates, as it resulted in extended lifespan of both sexes (female: z = -3.50, p < 0.001; male: z = -3.70, p < 0.001).

**Figure 2.**
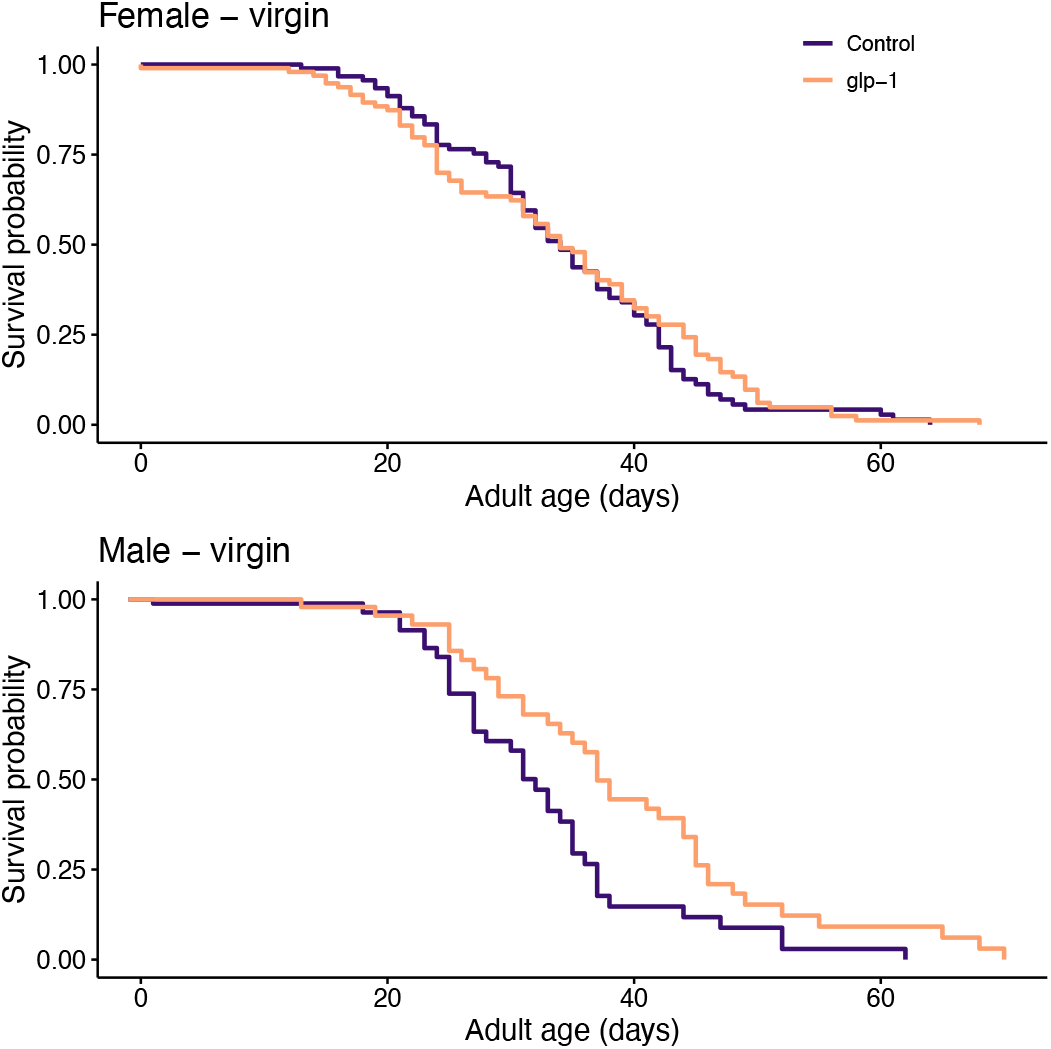
Virgin lifespan for females and males. Colour represent control (purple) or germline-less *glp-1* treated (orange) worms.

Germline removal resulted in a delayed development time to sexual maturity for females (χ^2^ = 9.26, d.f. = 1, p = 0.002), while the development time of males was not affected by germline removal (χ^2^ = 1.23, d.f. = 1, p = 0.268) (fig. 3A). Size at maturity was somewhat reduced (fig. 3B), although the effect on females was just non-significant (χ^2^ = 3.68, d.f. = 1, p = 0.055), while males significantly reduced size at maturity (χ^2^ = 4.83, d.f. = 1, p = 0.028). *Caenorhabditis* worms have most of their growth after maturity ^40^, but germline removal did not have the same effect on adult size for the two sexes (fig. 3C). For females, germline removal resulted in a substantially smaller adult size (χ^2^ = 17.25, d.f. = 1, p < 0.001), while adult size was not affected by germline removal in males (χ^2^ = 0.48, d.f. = 1, p = 0.487).

**Figure 3.**
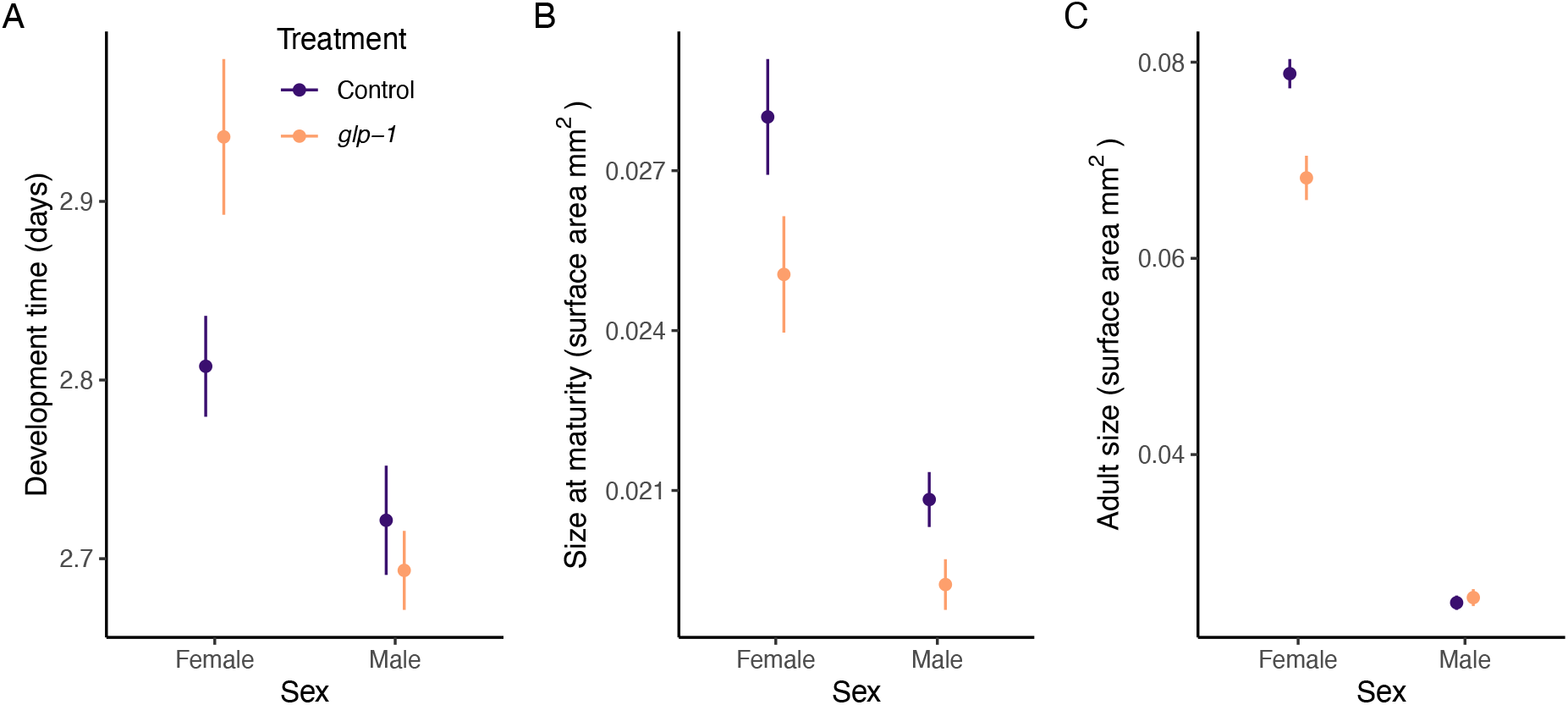
Development time (A), size at maturity (B) and adult size at day 4 of adulthood (C) for females and males. Colour represent control (purple) or germline-less *glp-1* treated (orange) worms. Error bars represent mean ± SE.

## Discussion

We found that germline removal using glp-1 RNAi results in increased heat-shock resistance in both sexes, but extended lifespan only in males. Since germline removal is predicted to free up resources previously allocated to germline maintenance ^11,23^, our findings suggest that germline maintenance could be more costly in males but the lifespan benefits may accumulate slowly over time.

There is good evidence for substantial costs associated with maintaining the germline and protecting it from mutations ^12,13^. For example, in a virus, there is a trade-off between replication fidelity and population growth ^43^, and in *Drosophila* high replication fidelity that evolved in a high UV treatment was partly lost under relaxed selection ^44^. Moreover, a recent study showed that male zebrafish exposed to radiation increase the repair of their germline at the cost of somatic repair. In contrast, males without a germline instead upregulate their somatic repair, providing compelling evidence for trade-off between somatic maintenance and germline maintenance ^23^. However, little is known about sex-specific effects of germline removal. While females invest relatively more in each gamete, males instead produce a large number of relatively small gametes ^24^. Consequently, males need to maintain a larger number of germline stem cells than females do. If maintenance of the germline is costly, males are expected to pay a higher cost of germline maintenance. This could be particularly pronounced in species like *C. remanei*, where males have longer reproductive lifespan than females ^40,45^ Our results support this hypothesis because germline removal increased male, but not female, lifespan.

Germline removal robustly increased heat-shock resistance in both sexes. Upregulation of heat-shock proteins have a protective effect against a range of stressors ^46^ and is generally seen as indicating investment in somatic maintenance ^47^. In *C. elegans*, heat-shock resistance is downregulated at the onset of reproduction and germline removal reduces the repressive chromatin marks on the heat-shock transcription factor HSF-1, which increase heat-shock resistance and lifespan ^48,49^. In *C. remanei*, high heat-shock resistance is associated with high condition ^36^, and the evolution of increased heat-shock resistance is costly, as it is traded off against investment in early-life traits ^37^. Our finding of increased heat-shock resistance in germline-less worms is thus in accordance with previous findings ^48,49^.

Selection for stress resistance often results in the evolution of long lifespan ^36,50^, long-lived lines are often stress resistant ^37^, and mutations in classic longevity genes are often increasing stress resistance ^51^. Therefore, both stress resistance and lifespan are often used as proxies for investment in somatic maintenance ^48^. Our finding that heat-shock resistance is increased by germline removal in both sexes, but lifespan only in males, suggest that the germline influences these traits partly independently. Indeed, stress resistance and longevity can be experimentally decoupled ^52^. Moreover, in *C. elegans* hermaphrodites, *glp-1* mutant worms, as well as worms subjected to dietary restriction, have extended lifespan and increased proteasome activity, suggesting a possible re-allocation of resources from germline to the soma through a stress response pathway that increase the proteasome activity ^53^. The increased proteasome activity in *glp-1* mutants is mediated by DAF-16/FOXO and results in upregulation of *rpn-6*.*1* (an essential subunit for the 26S/30S proteasome). However, while upregulated *rpn-6*.*1* activity increases heat stress resistance, it only increases lifespan under stressful conditions ^53^. Thus, lifespan and heat-shock resistance should be seen as related but distinct measures of somatic maintenance, and since germline-less *C. remanei* males increase both lifespan and heat-shock resistance, this suggests that germline maintenance may be more costly in males than in females.

Furthermore, germline removal reduced male growth before maturation but not in adulthood, while female growth rate was reduced both before and especially after maturation, where most growth occurs in *C. remanei* ^40^. Thus, germline removal improves male survival without any major growth costs, while germline-less females grow slower and do not live longer than reproductively functional counterparts in the absence of environmental stress. Body size is an important fitness-related trait in female nematodes ^40,54^ and a reduction in food amount ^55^ or quality ^56^ results in a reduction in body size. Since life-history traits such as lifespan and growth are often found to be negatively correlated ^6,37,57^, one could predict that females, instead of investing the freed up resources from germline removal in lifespan extension, would use if for increased growth. However, we find the opposite. Female *C. remanei* neither increase lifespan nor growth when the germline is removed, with the only benefit to somatic maintenance the females enjoy is increased heat-shock resistance. This could be either because fewer resources are being freed up by germline removal in females than in males, or because germline blocks the pro-longevity signalling only in males. These hypotheses are not mutually exclusive, and germline could potentially block pro-longevity signalling in males precisely because germline maintenance is costlier in males, who need to maintain functioning proliferative germline for longer in this species.

Interestingly, ablating the germline causes gigantism in *C. elegans*. This process is, unlike lifespan extension, independent of *daf-16* ^22^. The gigantism is, however, not caused by increased growth rate but by continued growth after day 4 of adulthood, when *C. elegans* hermaphrodites normally cease growth. We did not find that germline-less *C. remanei* were larger at day 4 of adulthood, they were instead smaller and therefore showing reduced growth rate. Although we did not measure size in late life, it is possible that they, like germline-less *C. elegans* hermaphrodites ^22^, may not cease to grow after day 4.

### Resource allocation or signalling trade-offs?

The expensive germline hypothesis postulates that the absence of a germline would free up resources from germline maintenance that could instead be used for somatic maintenance ^11^. Specifically, this hypothesis predicts that germline removal should extend lifespan in non-reproducing organisms and costs should be more pronounced in males who need to maintain larger number of germline stem cells across life course ^11^. Our results provide at least partial support as germline removal increased heat-shock survival in both sexes compared to virgin controls, but increased lifespan only in males.

However, we also documented increased development time to maturity, reduced growth rate and smaller adult body size in females suggesting the germline removal has strong sex-specific effects on the whole organism life-history. This poses the question if the sex-specific lifespan extension following germline removal is a consequence of more resources being freed up for somatic maintenance in germline-less males, or a result of sex-specific regulation of lifespan by the germline signalling? It is likely that resistance to heat-shock is beneficial to *C. remanei* in their natural environment, and germline-mediated reduction in resistance could reduce fitness of an adult organism. However, it has been shown in *C. elegans* that resistance to heat-shock is downregulated in adult organisms following maturation ^47^. While this could be a result of resource allocation trade-off, it can also result from insufficient selection on late-life performance ^29^, especially because downregulation of heat-shock resistance occurs in *ad libitum* food and in a benign environment. Ultimately, future studies should aim to directly test for increased use of resources by somatic tissues following germline removal in both sexes.

### Cost of mating

Notably, the survival benefit of germline-less males was only present if the worms were virgin. If mated, germline removal did not extend lifespan in any sex, even though neither sex produced any offspring. This suggests that high costs of mating strongly reduce lifespan in both sexes. In *C. elegans*, mating reduces lifespan in germline-less (*glp-1*) hermaphrodites ^58^ and even the presence of males reduces lifespan of hermaphrodites through secreted compounds ^59^. Our findings that the effect of germline removal on lifespan was only detectable in virgin worms mirror a previous finding on the effect of rapamycin in *C. remanei*, where its effect on lifespan was stronger in virgin than in mated worms ^40^.

### Summary

In summary, we found that germline removal increased heat-shock resistance of both sexes, but lifespan only in males. Moreover, females paid a cost of substantially reduced adult size, while male adult size was not affected. Together, these results suggest that germline differentially regulates male and female life-histories. The results are consistent with the hypothesis that germline maintenance is more costly for males than females, and this may be a substantial, but understudied, cost of reproduction in males ^60^.

## Author contribution

MIL, BSM and AAM designed the experiment; AH, HC and EG performed the injections, BSM, HC and MIL performed the phenotypic assays, MIL analysed the data, MIL drafted the manuscript together with AAM. All authors contributed to the revision of the manuscript and gave final approval for publication.

## Acknowledgements

The authors are grateful to Dr. A. Liontos and Dr. Y. Zhao for technical assistance with RNAi constructs.

## Supplementary materials

**Figure S1.**
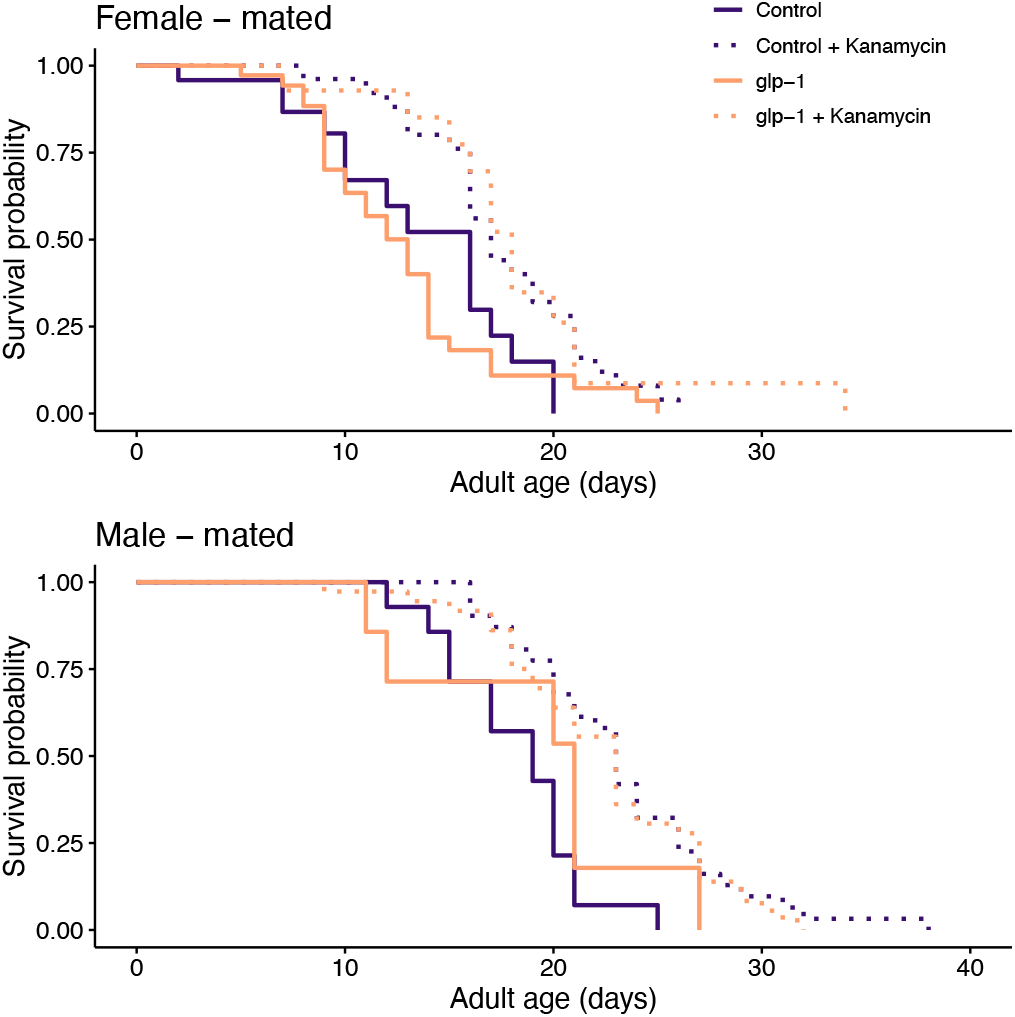
Mated lifespan for females and males. Colour represent control (purple) or germline-less *glp-1* treated (orange) worms. No effect of germline removal on lifespan was detected in any sex.

